# Selection of different parameters to study epistatic effect between heat shock and slightly deleterious mutations in the most fragile stage of embryogenesis *Cyprinus carpio* L

**DOI:** 10.1101/2024.01.23.576732

**Authors:** Aleksandr Kuzmin, Viktoria Skripskaya, Valeria Lobanova, Polina Brodovaya, Alina Tarasova, Konstantin Popadin, Alina G. Mikhailova, Elena Kozenkova, Nikolai Mugue, Evgenii Vinogradov, Dmitrii Balashov

**Affiliations:** Center For Mitochondrial Functional Genomics, Institute of Living Systems, Immanuel Kant Baltic Federal University, Russia; Branch for the freshwater fisheries of the Federal State Budget Scientific Institution “Russian Federal Research Institute of Fisheries and oceanography” Laboratory of genetics and fish; Federal State Budget Scientific Institution “Russian Federal Research Institute of Fisheries and oceanography” Laboratory of genetics and fish breeding (VNIRO); Moscow Institute of Physics and Technology (MIPT), Moscow, Russia

**Keywords:** purifying selection, deleterious mutations, carps, heat shock

## Abstract

The primary genetic challenge encountered in artificial populations lies in the strong genetic drift, which leads to the accumulation of numerous slightly deleterious mutations across the genome. Such mutations diminish the adaptability of the entire population. The objective of this project involves the investigation and implementation of genetic selection methods within cultivated fish populations such *Cyprinus carpio L*. In order to maintain a high level of genome quality in productive species, we conducted a proof-of-principle experiment employing stress-induced strong purifying selection. This selection process is based on negative epistasis and effectively eliminates organisms carrying an excess of deleterious variants. The first step involves the creation of mutant and intact groups of fish. To obtain mutant groups, we treated male gametes with the ENU mutagen, which primarily induces single-nucleotide substitutions uniformly throughout the genome, thereby imitating natural mutations. This methodology is paramount for the accurate interpretation of experimental outcomes. Notably, temperature stands as a pivotal factor influencing the embryonic development of fish. Therefore, we subjected the embryos to a diverse range of temperatures and varied the duration of exposure during critical stages of embryogenesis. Through meticulous examination, we ascertained that the stage most susceptible to screening purposes is the 22-somite pair stage, occurring at a temperature of 38°C, with a 40-minute exposure period. We suppose, this comprehensive approach can be applied to improve the quality of the gene pool within domestic fish populations, ultimately enhancing the economic efficacy of fish farms. The future prospects of this method encompass its potential application to various species.

## Introduction

The genetic underpinnings of artificial populations are critical to their viability and productivity. Genetic drift in these populations can lead to an accumulation of slightly deleterious mutations, which threatens their adaptability and long-term sustainability^1,2^. This phenomenon is not only a concern in the realm of aquaculture but is also a well-documented consequence of domestication and improvement in a variety of species, from plants to animals^3–7^.

In the context of *Cyprinus carpio L*., a species of significant aquacultural value, the objective of our project is to investigate and implement genetic selection methods to counteract these genetic challenges. We approach this by conducting a proof-of-principle experiment that utilizes stress-induced purifying selection to remove organisms with an excess of deleterious mutations, a method supported by recent findings on negative epistasis and its role in maintaining genetic health^8,9^.

The experimental protocol begins with the controlled induction of mutations in male gametes using ENU mutagen, thereby emulating the natural mutation process^10,11^. This controlled induction is critical to understanding the mutational landscape and sets the stage for subsequent selection processes.

Temperature is a critical abiotic factor that significantly influences development of fish embryos. The necessary thermal window for the induction of sexual maturation, oogenesis, and viability of offspring fluctuates from 19°C to 30°C, with an optimal median at approximately 23°C^12^. Deviations from this thermal optimum induce the biosynthesis of Heat Shock Proteins (HSPs), a cytoprotective response facilitating organismal survival under suboptimal thermal conditions^13,14^. Individuals burdened with an accumulation of slightly deleterious mutations exhibit a reduced probability of endurance of extreme environmental stressors. This reduction in viability can be attributed to the inability of molecular chaperones to effectively mitigate the amplified mutational load in conjunction with adverse environmental conditions. Consequently, this selection pressure favors the persistence of genotypes characterized by genetic resilience. It is for this reason that thermal regulation has been selected as the environmental stressor of choice for conducting purifying selection within our research aim.

The fertilization stage is succeeded by the pharyngula period, during which the embryo undergoes significant morphogenesis, culminating in the “eye stage,” a crucial juncture for the application of temperature shock^15^.

The implications of this study are manifold, extending beyond the immediate improvement of *Cyprinus carpio L*. populations to broader applications in the genetic conservation and management of domesticated species. The genetic foundation laid down by domestication, as reported in various species, indicates a shared burden of genetic costs that our methodology aims to alleviate^16–18^.

## Materials and Methods

### Preparations

#### Selection and capture of carp producers and their subsequent preparation for obtaining reproductive products

After winter, carp producers are carefully transferred to designated pre-spawning housing, where males and females are kept separately^19^. As the water in the ponds gradually warms up and reaches optimal spawning temperatures, the fish are then moved to trays within the hatchery workshop. Here, the factory method of artificial reproduction is employed, allowing for controlled fertilization and the collection of reproductive products^19^. This meticulous process ensures the efficient propagation of carp populations and contributes to the sustainable management of fish stocks.

#### Initiation of gamete maturation in carp producers

Females undergo an injection of a suspension of acetonated carp pituitary glands at a dosage of 4 mg/kg (total dose), which triggers ovulation. This injection is administered in two stages: a preliminary injection of 10% of the total dose is given to the females 24 hours prior to egg collection, followed by another injection after 12 hours. Males, on the other hand, receive half the dosage of the hormone over a 24-hour period, with a pre-injection into the females. Throughout this process, the producers are housed in trays that maintain optimal temperature and oxygen conditions. After 12 hours from the final injection, eggs and sperm are obtained, ensuring the availability of reproductive materials for artificial reproduction purposes.

#### Obtaining reproductive products from carp

To collect fish sperm for fertilization, a designated room is prepared with dry and clean containers. Prior to obtaining the sexual products, the fish are gently dried using a soft towel, ensuring that the genital opening is wiped dry with gauze. The collected sperm is then analyzed under a microscope, assessing its motility, which should be at least 90% forward movement^20,21^.

When it comes to obtaining caviar from female carp, clean and dry enamel basins are used. It is crucial to prevent any water from entering the basins, either from the fish or from the hands of the fish farmer. Similarly, prior to obtaining the sexual products, the fish are carefully dried with a soft towel, and the genital opening is wiped dry with gauze. The eggs of each female are collected in individual containers. Normally, ovulated eggs are released with gentle pressure on the abdomen, moving from head to tail. This meticulous process ensures the collection of viable reproductive materials for artificial reproduction purposes.

#### Fertilization and incubation of carp embryos

Fertilization takes place in controlled settings using plastic Petri dishes with a diameter of 100 mm. A clean and dry cup is prepared, and a portion of caviar containing approximately 50-100 eggs is added. Next to the caviar, a drop of sperm (10 μl) is placed, followed by the addition of water (20-40 ml) to the cup. The sexual products are gently mixed together using a bird’s feather. Fertilization occurs within a duration of 60 seconds, after which the activated eggs adhere to the bottom of the Petri dish. The water used for fertilization is then drained, and freshwater (50-60 ml) is added.

Incubation of fertilized eggs occurs at a constant temperature of 20°C. The following day, any dead unfertilized eggs are removed, and the percentage of fertilization is calculated. It is crucial to replace the water in the cups with fresh water twice a day. Incubation should be conducted in a dark or shaded area, ensuring the exclusion of direct sunlight. These controlled conditions optimize the development and survival of carp embryos during the incubation process.

### Heat shock preparation

To induce heat shock, it is imperative to employ a thermostatically regulated container, specifically a bath with a volume of approximately 20 liters. This bath should possess the capability to accommodate 30 Petri dishes simultaneously. It is essential that the bath be equipped with heating elements, a temperature measuring device, and a thermostat. In order to sustain the ideal environment for the shock process, adequate water circulation must be achieved through the utilization of a pump, coupled with aeration mechanisms.

### N-ethyl-N-nitrosourea mutagenesis

For the preparation of a 2 mM solution of N-ethyl-N-nitrosourea (ENU), 0.0324 g of ENU is dissolved in 1.5 mL of a 10 mM sodium acetate. Subsequently, 3.65 mL of a 0.1M Hanks salt solution and 1.25 mL of sperm are combined. Then 100 µL of ENU solution is added into the Hanks salt solution containing the sperm. After 45 min of sperm incubation wild-type eggs were fertilized as described above.

## Results

### 1. Selection of heat shock modes (Experiment 1 2022)

To create conditions for thermal shock, we utilized a short-term increase in ambient temperature with various exposure regimes in order to determine the semi-lethal dose eliminating 50% (LD50) of individuals (eggs). Based on literature sources^22^, we introduced four thermal regimes: 30, 32, 34, 36°C and 20°C as control, along with exposure times of 10, 15, 20, and 25 minutes.

Based on the results of the initial experiment, we did not observe a significant effect of the induced heat shock (i.e., egg mortality) (Figure 1). Insufficient temperature impact on the eggs was noted. We attribute the lack of impact to well-developed embryonic membranes, which negate any temperature stress. As a result, we developed two strategies to determine the LD50 in these subjects: (i) a sharp increase in the temperature regime, which is likely to have a negative effect on embryonic development, and (ii) prolonging the exposure time alongside a slight increase in the temperature regime.

**Figure 1.**
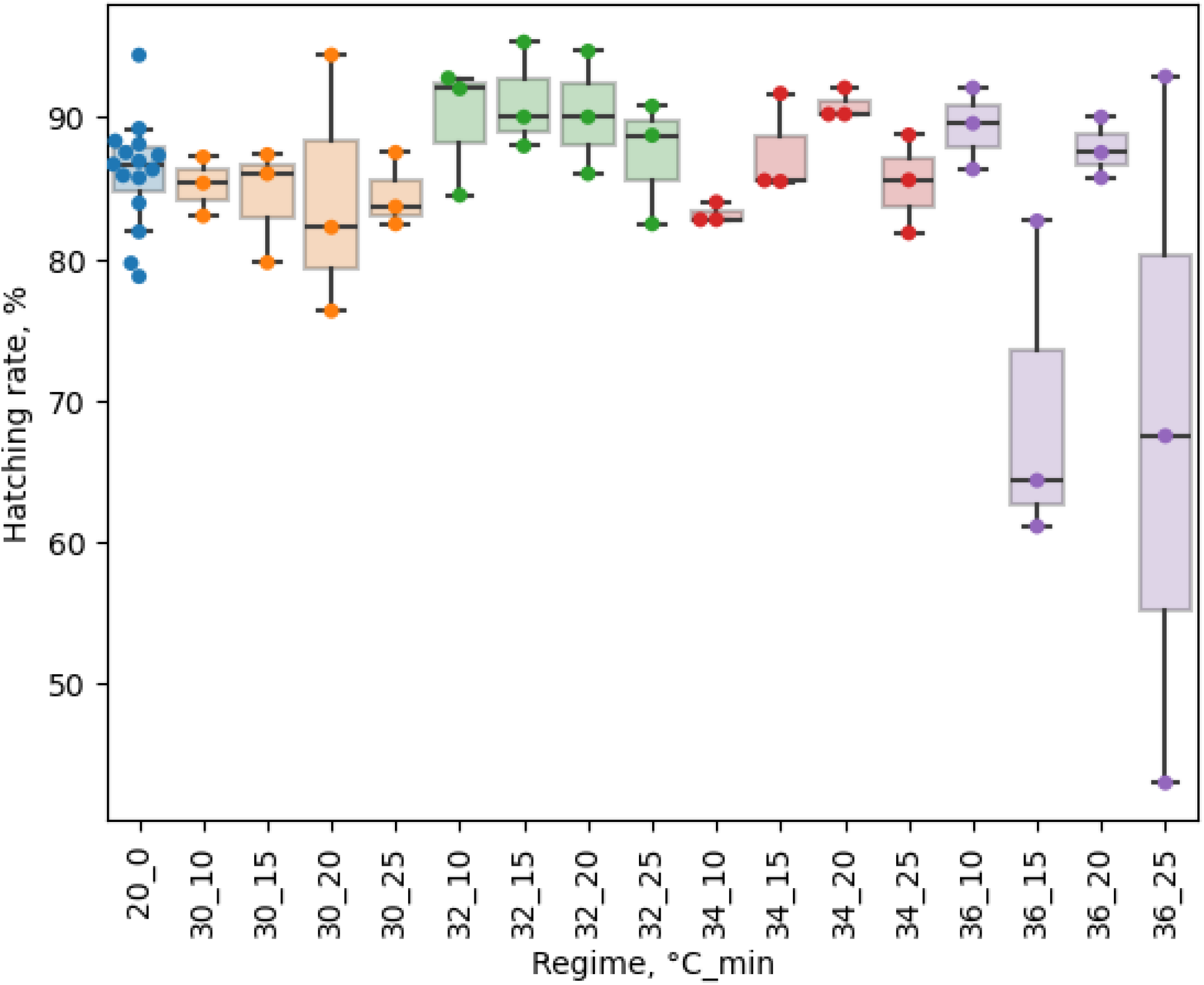
Distribution of Hatching rate among different regimes. Regimes are no different statistically significantly from the control (U-test, Bonferroni correction).

### 2. Selection of heat shock parameters with induced mutagenesis (Experiment 2 2022)

During the experiment, the fertilization of carp eggs was conducted using sperm diluted at an ENU (Ethyl Nitrosourea) concentration of 0.55 mM, as described^10^. The sperm dilution was prepared using solutions of sodium acetate (17 mM) and Hanks salts (9.8 mg/mL). This resulted in two groups of carp eggs: one group fertilized with native (non-mutagenic) sperm and another group fertilized with mutagenic sperm containing ENU.

Following fertilization, heat shock parameters based on a previous experiment (selection heat shock modes) were applied to select the LD50 based on different temperature regimes and exposure times. The objective was to increase the exposure time and temperature in order to surpass the homeostatic barrier of the eggs. The following temperature regimes were employed on the two groups of carp eggs: 37°C for 30 minutes, 37°C for 60 minutes, 40°C for 30 minutes, 40°C for 60 minutes, and 20°C used as a control. After the 24-hour period, the number of fertilized eggs and dead eggs were counted and recorded. Another day later, the hatching and swimming abilities of the remaining eggs were assessed.

The results indicate that none of the tested regimes exhibited statistically significant effects on the fertilization rate (U-test with Bonferroni correction) (Figure 2). However, the temperature regimes of 40°C for 30 minutes and 40°C for 60 minutes displayed a significant impact on the hatching rate. These findings suggest that the applied temperature regimes were wide-ranging, with 37°C proving insufficient for achieving the desired heat shock parameters, while 40°C proved to be excessively high for the eggs.

**Figure 2.**
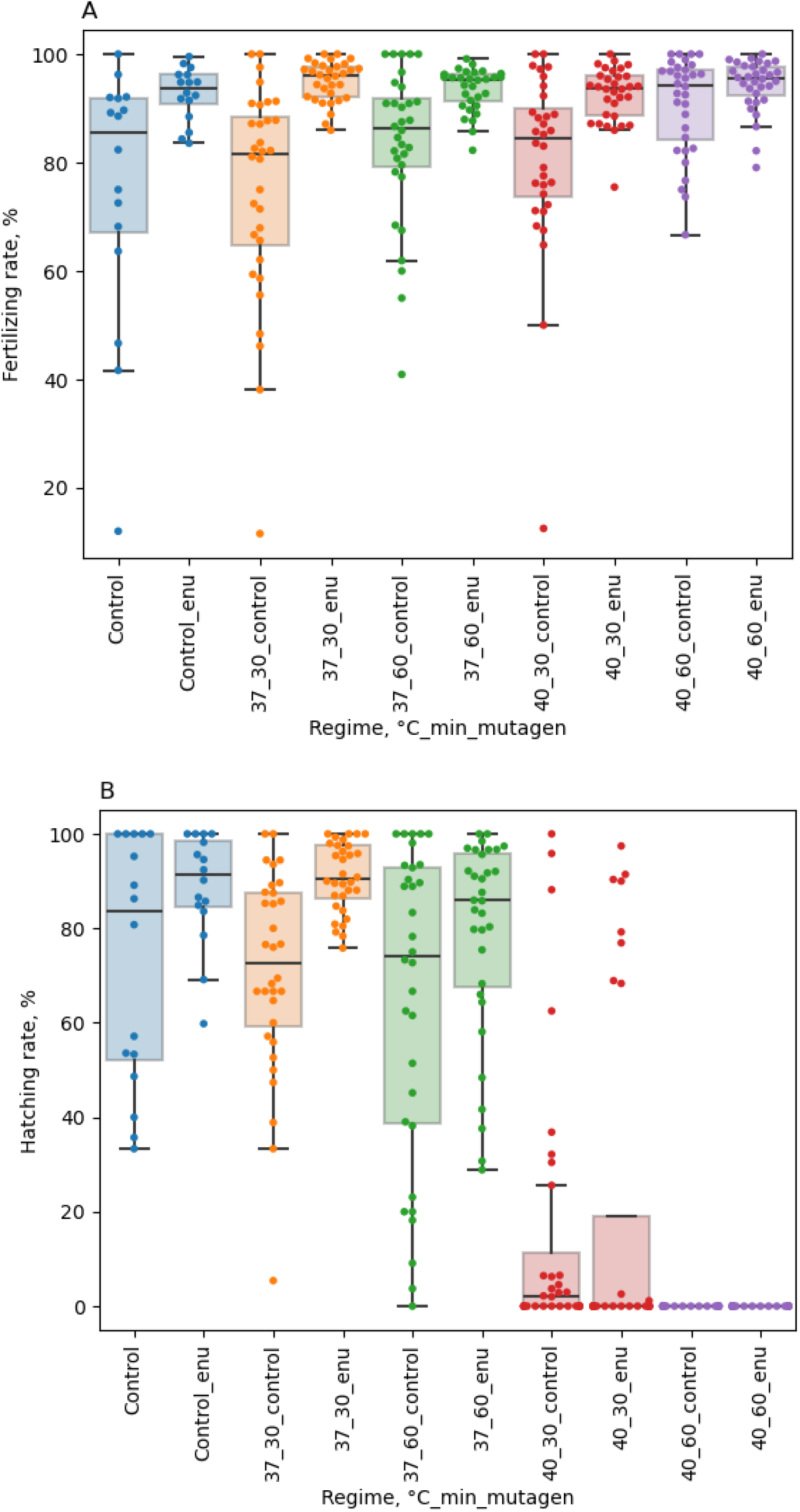
Distributions of Fertilizing rate (A) and Hatching rate (B) for different heat shock regimes with induced mutagenesis. All regimes are no different statistically significantly from the control (U-test, Bonferroni correction) except 40°C for 30 minutes and 40°C for 60 minutes. The median hatching rate in these two groups is 2% and 0% respectively.

### 3. Selection of mutagen concentration (Experiment 3 2022)

In order to determine the LD50 (lethal dose for 50% of the population) of ENU on carp eggs, the following experimental procedure was conducted. Carp eggs were first fertilized with sperm that was diluted with various concentrations of ENU. The sperm dilution was prepared using solutions of sodium acetate (17 mM) and Hanks salts (9.8 mg/mL). The concentrations of ENU used were 0 mM, 1 mM, 2.5 mM, 5 mM, and 10 mM.

To initiate fertilization, approximately 80-100 mg of carp caviar was added to petri dishes using a spoon. Subsequently, a volume of 1.25 mL of the ENU-diluted sperm was added to the petri dishes. The fertilization process was then carried out according to the methods. After 10 minutes of exposure, the water in the petri dishes was replaced with fresh water. This step aimed to remove any unfertilized eggs and residues of the mutagen (ENU). The petri dishes were then placed in a thermostat set at a temperature of 17.5 °C and left undisturbed for 24 hours. After the 24-hour period, the number of fertilized eggs and dead eggs were counted and recorded. Another day later, the hatching and swimming abilities of the remaining eggs were assessed.

Based on the data analysis, it was determined that ENU concentrations exceeding 5 mM have a lethal effect on fish development (Figure 3). Consequently, for subsequent experiments, the concentrations of 1.5 mM and 2 mM were chosen to minimize the mortality rate and allow for the evaluation of other developmental parameters.

**Figure 3.**
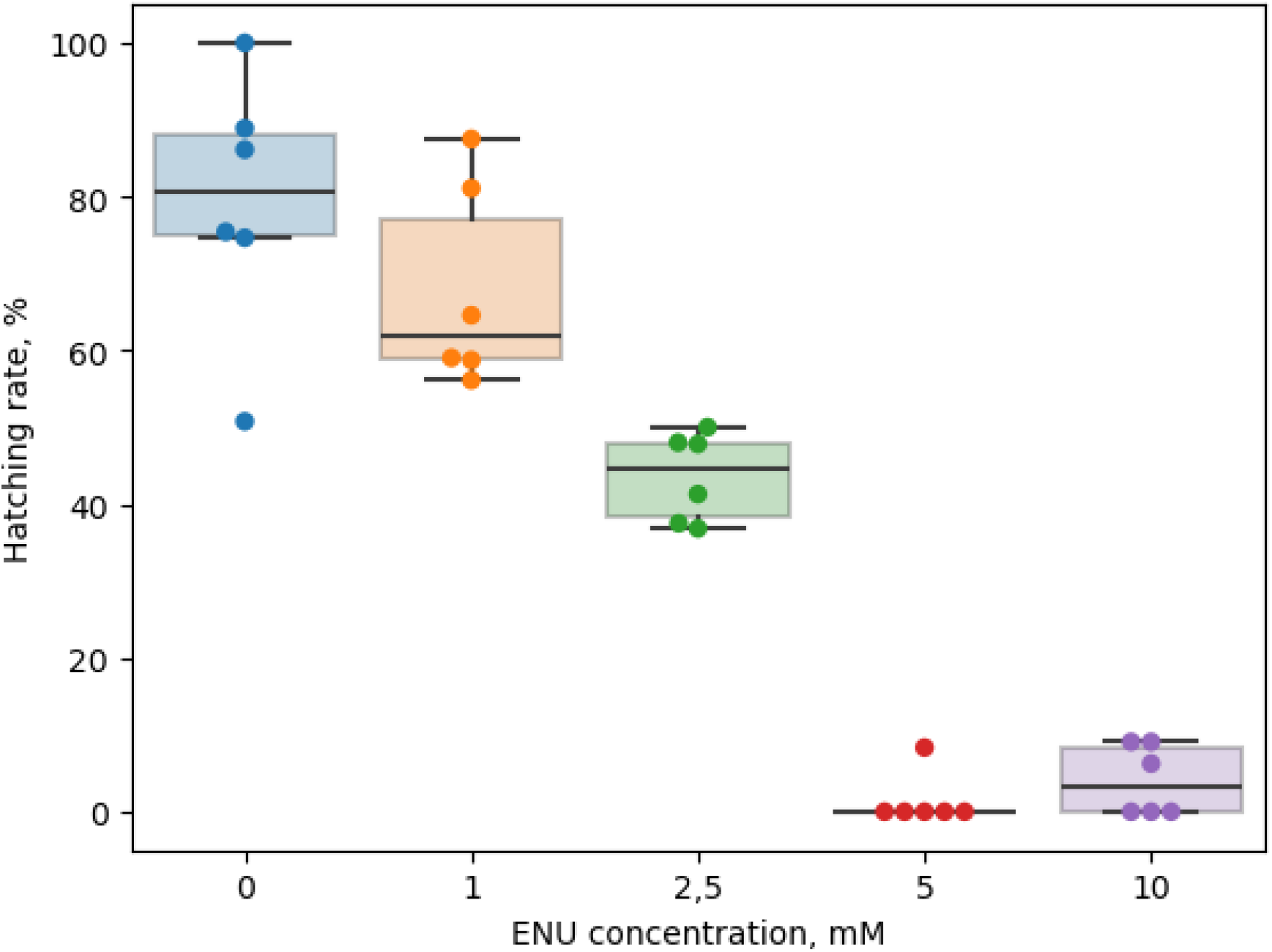
Distributions of Hatching rate for various mutagen concentrations. The group with an ENU concentration of 1 mM shows no statistically significant difference when compared to the control group. Conversely, Group 2,5 demonstrates significant statistical differences in comparison to the other groups. Additionally, Groups 5 and 10 do not display any statistically significant differences between each other; nevertheless, they do exhibit significant differences when compared to the remaining groups (ANOVA, Tukey’s HSD).

**Figure 4.**
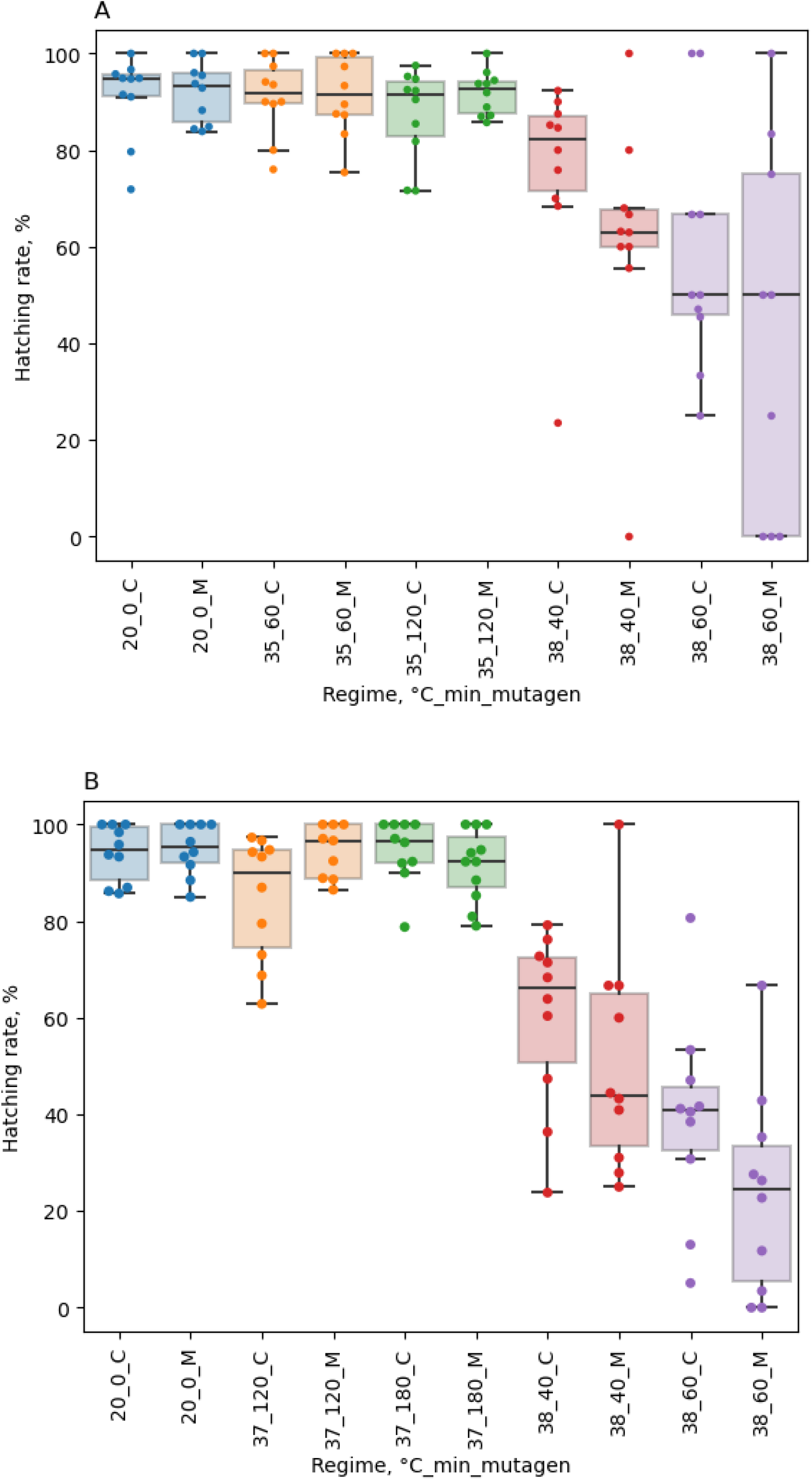
Distributions of Hatching rate for (A) 22 somite stage, (B) pharyngula stage. Regimes with a temperature of 38°C demonstrate a statistically significant divergence from the control group (U-test, Bonferroni correction).

**Figure 5.**
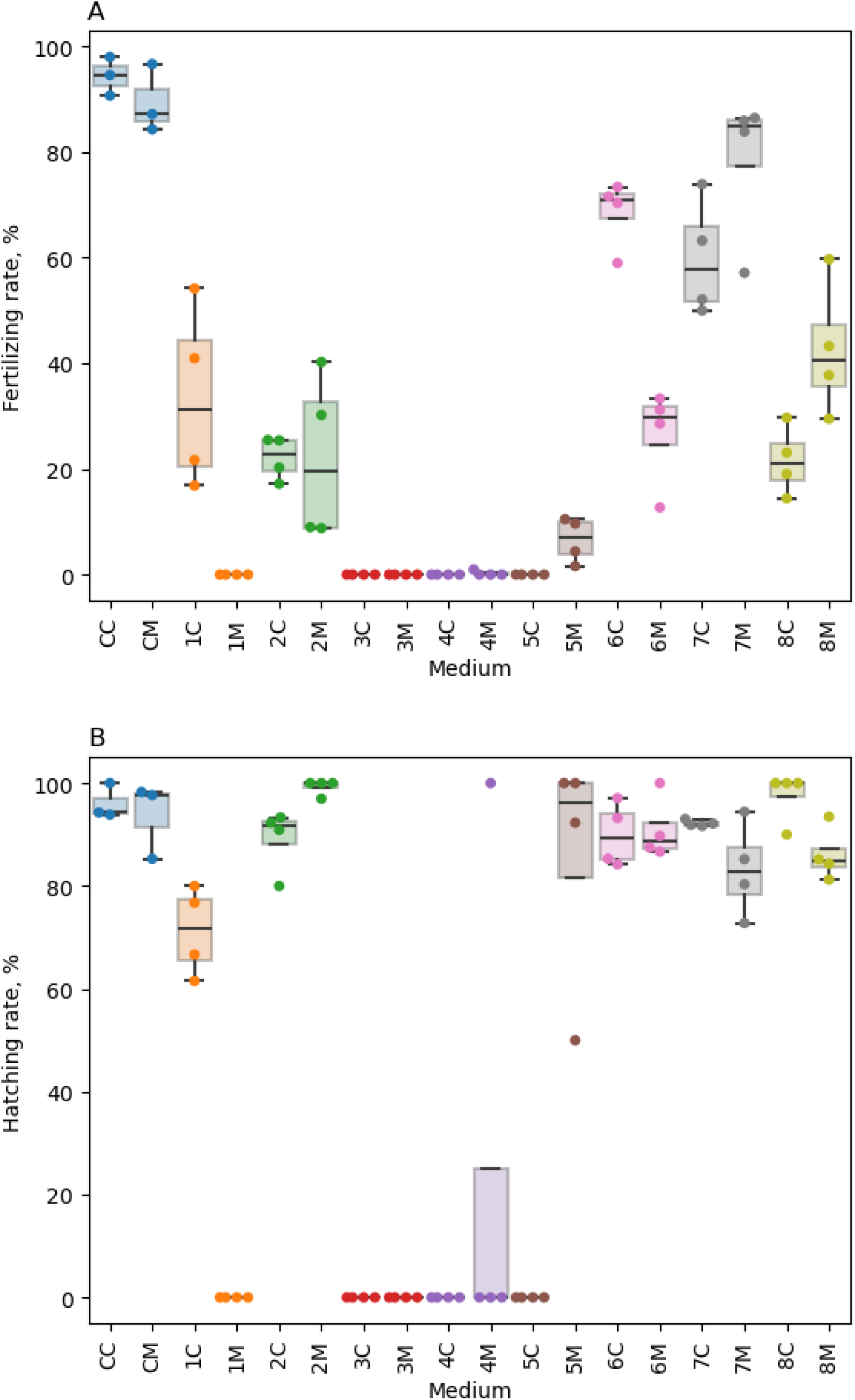
Distributions of Fertilizing rate (A) and Hatching rate (B) for various mediums. CC - not frozen not mutant sperm, CM - not frozen mutant sperm, 1C - frozen not mutant sperm, 1M - frozen mutant sperm, 2C - frozen sperm in medium 2, 2M - frozen mutant sperm in medium 2, 3C - frozen sperm in medium 3, 3M - frozen mutant sperm in medium 3, 4C - frozen sperm in medium 4, 4M - frozen mutant sperm in medium 4, 5C - frozen sperm in medium 5, 5M - frozen mutant sperm in medium 5, 6C - frozen sperm in medium 6, 6M - frozen mutant sperm in medium 6, 7C - frozen sperm in medium 7, 7M - frozen mutant sperm in medium 7, 8C - frozen sperm in medium 8, 8M - frozen mutant sperm in medium 8.

### 4. Selection of stage of embryonic development stage for heat shock application with induced mutagenesis (Experiment 1 and 2 2023)

In this experiment, we aimed to explore the impact of heat shock at different stages of embryonic development. Heat shock treatments were applied to carp embryos at the gastrula, 22 somite stage, and pharyngula (pigmentation) stages. The fertilization of carp eggs was carried out using sperm diluted at an ENU concentration of 2 mM, following the same protocol as mentioned earlier. As a result, two groups of carp eggs were obtained: one fertilized with native sperm and the other fertilized with mutagenic sperm containing ENU. The heat shock parameters used in this experiment were based on previous studies, where temperatures of 35°C, 37°C, 38°C, and a control temperature of 20°C were employed. The exposure time to heat shock varied between 30 minutes, 40 minutes, 60 minutes, 120 minutes, and 180 minutes.

Upon analyzing the results, it was observed that the gastrula stage proved to be highly vulnerable to heat shock with induced mutagenesis. In fact, the majority of fertilized eggs, irrespective of whether they were fertilized with native or mutagenic sperm, experienced significant mortality following the heat shock treatment. This indicates that the gastrula stage is too fragile to withstand the detrimental effects of heat shock, making it unsuitable for experimental manipulations involving induced mutagenesis.

Conversely, the pharyngula stage appeared to be less susceptible to the detrimental effects of heat shock. The embryos at this stage displayed significantly higher viability and demonstrated minimal susceptibility to the detrimental effects of heat shock. Furthermore, both the native and mutagenic fertilized eggs at the pharyngula stage exhibited robust survival rates, indicating that this stage is relatively unaffected by heat shock-induced mutagenesis. These findings suggest that the pharyngula stage is a resilient developmental period that is also unsuitable for experimental manipulations.

Interestingly, the 22 somite stage demonstrated to be the least appropriate stage for conducting experiments involving heat shock. The embryos at this stage exhibited lower viability and resilience to the heat shock treatment compared to the other stages tested.

## Conclusions

In conclusion, this study has underscored the criticality of managing genetic quality within artificial populations, with a focus on *Cyprinus carpio L*. By employing a novel approach to purifying selection through the stress-induced method, we have demonstrated significance of mitigating genetic drift. The targeted use of ENU on male gametes and application of heat shock during the 22-somite pair stage of embryogenesis have proven effectiveness of heat shock in enhancing the genetic robustness of carp populations. The effectiveness of such selection methods could help with genetic management in fish farms, potentially leading to greater economic benefits and a sustainable future for domestic animal populations. Additionally, we tested the effect of mutagen and cryoprotectors on sperm quality and haven’t observed any deleterious effects (Supplementary Material).

## Acknowledgments

AK and KP were supported by the Ministry of Science and Higher Education of the Russian Federation (agreement no. 075-15-2021-1084) for design of the experiment, performance of the experiment and analysis of the results.

