## Supplementary Material for "Selection of different parameters to study epistatic effect between heat shock and slightly deleterious mutations in the most fragile stage of embryogenesis *Cyprinus carpio* L"

### Effect of mutagen on sperm quality (Experiment 6 2023)

We also investigate the effect of mutagen, cryoprotector (CP) and freezing on sperm quality and the ability to fertilize native eggs. The cryoprotector consisted of 20% methanol and 6% ethylene glycol by weight.

The fertilization of carp eggs was carried out using sperm diluted at an ENU concentration of 2 mM, following the same protocol as mentioned above. As a result, two groups of carp eggs were obtained: one fertilized with native sperm and the other fertilized with mutagenic sperm containing ENU. Next we added 8 different mediums (Supplementary Table 1) to sperm, immediately frozen on -80. Then unfroze, checked sperm quality (sperm motility), and fertilized wild-type eggs.

Motility of native sperm was 86%.

Supplementary Table 1 - Composition of media

| № | Sperm dilution (v/v) | Diluent or medium, order of adding CP |
|---|---|---|
| 1C | 1:3 | sperm + HBSS without CP + 20% methanol, 6% ethylene glycol |
| 2C | 1:3 | sperm + HBSS with 20% methanol, 6% ethylene glycol |
| 3C | 1:3 | sperm + 0,1% sucrose solution, 0,3% KCl + 20% methanol, 6% ethylene glycol |
| 4C | 1:3 | sperm + 0,1% sucrose solution, 0,3% KCl, 20% methanol, 6% ethylene glycol |
| 5C | 1:3 | sperm + 0,1% sucrose solution, 0,35% NaCl + 20% methanol, 6% ethylene glycol |
| 6C | 1:3 | sperm + 0,1% sucrose solution, 0,35% NaCl, 20% methanol, 6% ethylene glycol |
| 7C | 1:1 | sperm + 0,1% sucrose solution, 0,35% NaCl, 20% methanol, 6% ethylene glycol |
| 8C | 1:2 | sperm + 0,1% sucrose solution, 0,35% NaCl, 20% methanol, 6% ethylene glycol |
| 1M-8M | | same with mutagen |

Dokina O.B., Kovalev K.V., Pronina N.D., Milenko V.A., Kovalenko V.N. Optimized technologies for large-scale cryopreservation of carp fish sperm // Fishery. - 2022. - № 6. - C. 58-66
